# Paradoxical influences of prediction are resolved across time

**DOI:** 10.64898/2026.03.05.709935

**Authors:** Kirsten Rittershofer, Yuyang Wang, Martin Eimer, Peter Kok, Daniel Yon, Clare Press

## Abstract

It is widely thought that our brains use expectations to optimise perception, yet theoretical accounts disagree about the form of this optimisation. Bayesian accounts propose that perception is biased towards expected events, rapidly generating more veridical experiences, whereas cancellation accounts argue that unexpected inputs are perceptually prioritised as they are more informative. A recent opposing process theory reconciles these views by proposing a temporal reversal in prioritisation: perceptual processing is pre-emptively biased towards what is expected, followed by later enhancement of only particularly surprising inputs that are informative for learning and model updating. Here, we tested this account using time-resolved decoding of EEG data, while participants observed avatar action outcomes that were either expected or unexpected based on their own movements. Expectation effects on the neural representations of these action outcomes indeed unfolded over time in such a manner, with higher decoding for task-relevant expected outcomes starting before stimulus onset, followed by a later post-stimulus advantage for unexpected outcomes. These findings support the opposing process theory, demonstrating a reversal in prioritisation that is not predicted by current Bayesian or cancellation accounts. By exerting distinct influences across time, expectations can thus render perception veridical, while still allowing for the accurate perception of particularly unexpected events that are important for updating our beliefs or ongoing courses of action.

**Significance statement:** How we perceive the world is shaped by our expectations. Yet it remains unclear how the brain can use these expectations to make perception both more accurate and more informative, given these demands require opposite influences of expectation on processing – upweighting versus downweighting the expected, respectively. Here, we show that this apparent conflict is resolved across time. The expected is enhanced before a stimulus appears by pre-activating what we expect, whereas unexpected inputs are enhanced later. This temporal reversal allows expectations to support both demands, rapidly generating generally more accurate perceptual experiences, while also remaining sensitive to unexpected information that is critical for learning and adaptive behaviour.

## Introduction

Our sensory systems are constantly bombarded with noisy input and it has been widely suggested that our brains must thus rely on expectations in order to perceive (1–5). However, it remains unclear *how* expectations shape perception. Bayesian accounts propose that perception is biased towards what we expect (6–8). Such a mechanism would generate more veridical experiences as expected events are, by definition, more likely to occur, while also compensating for processing delays by enabling best guesses to be generated even prior to stimulus onset. Supporting this view, numerous studies show that perception is biased towards expected events (9–11), which reach conscious awareness more rapidly (12–15) and are perceived more clearly and intensely (16–18). In contrast, cancellation accounts argue that unexpected events are perceptually prioritised, as these are more informative. That is, they tell us something we did not already know, and are hence important for learning, model updating, and action planning (19,20). These accounts were initially conceptualised to explain action perception and are hence supported by studies reporting that expected action consequences are perceived less intensely (21–25). More recently, similar findings have been reported across sensory domains, suggesting that unexpected events are prioritised in a number of ways (26,27). At the neural level, findings are likewise mixed, with expectation effects observed in opposing directions across domains and imaging modalities (28–35).

Given these two conflicting accounts, each supported by a host of empirical evidence, the question arises of how this apparent paradox might be resolved. That is, it should be adaptive to use expectations to render perception veridical, fast, *and* informative, rather than trading one adaptive benefit for another. The opposing process theory offers a potential solution, by outlining how expectations can support all of these demands (36). According to this account, perception is initially biased towards what we expect, rapidly generating largely veridical percepts. However, when inputs strongly deviate from expectations, a later reactive process selectively enhances the perception of these highly surprising events, as they are likely informative for learning and model updating, while ignoring smaller deviations likely due to noise.

In the present study, we tested this account using expectations about action outcomes. Participants performed finger movements and observed an onscreen avatar hand synchronously performing either the expected (matching) or unexpected (mismatching) action. We orthogonally manipulated the task relevance of the observed action outcomes to determine whether any effects were modulated by attention. To assess the predicted temporal dynamics, we used multivariate decoding of electroencephalography (EEG) signals to track the quality of the neural action outcome representations with high temporal resolution. While the paradox concerns perceptual experience, the temporal dynamics are proposed to be observable in neural activity. Specifically, the opposing process theory predicts enhanced neural representations (higher decoding) for expected outcomes early in time, potentially even before stimulus onset due to pre-activation of the expected sensory outcome representation (31,37), followed by later enhancement of unexpected outcomes. In contrast, the other two accounts do not predict such a temporal reversal, but rather consistently higher decoding for either expected (Bayesian accounts) or unexpected outcomes (cancellation accounts; Figure 1D).

**Figure 1:**
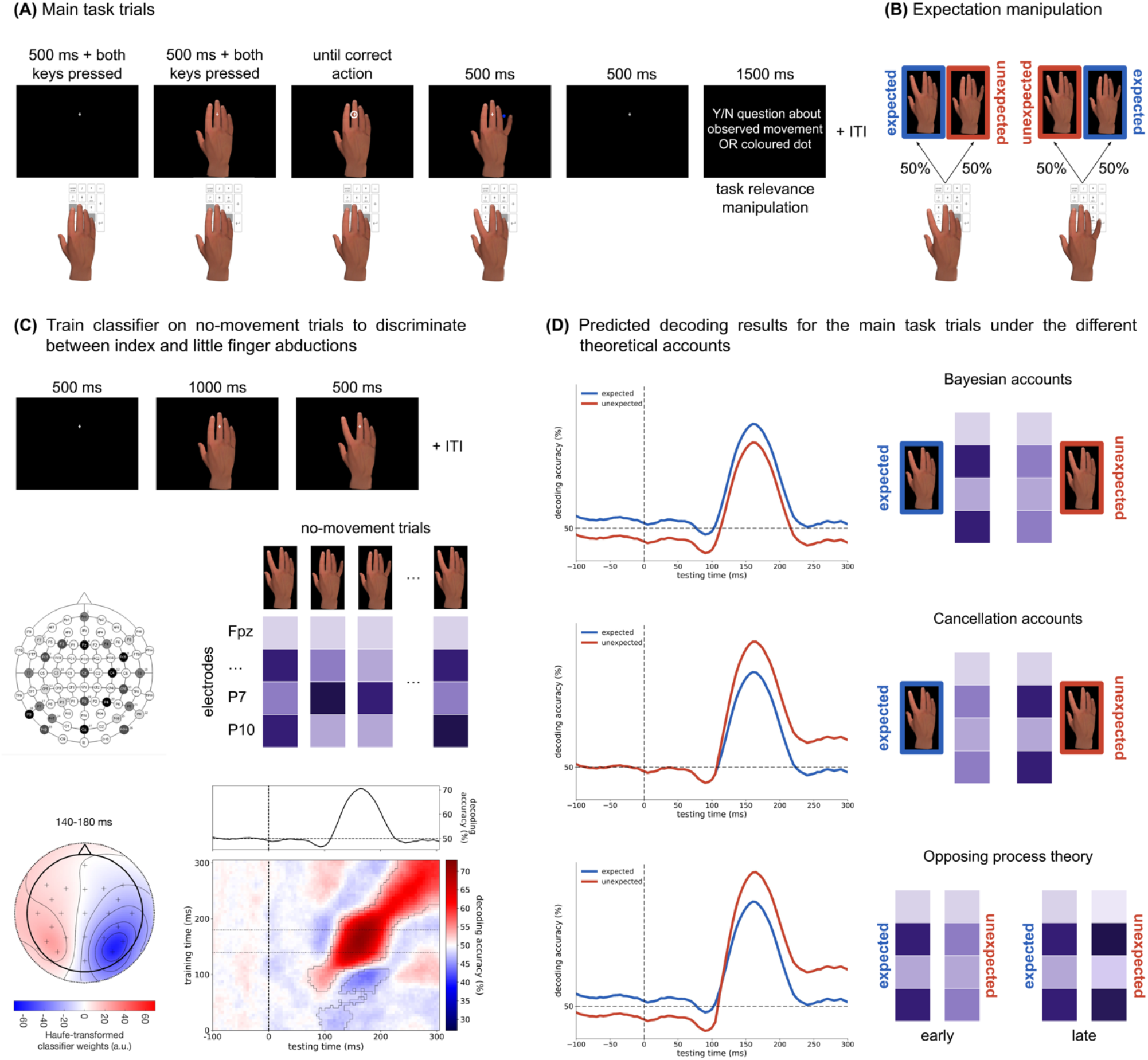
Experimental design and predicted results under the different theoretical accounts. **(A)** Main task. Participants abducted either the index or little finger of their right hand in response to a shape cue (here: circle = index finger). In synchrony with the participant’s action, an avatar hand on the screen moved. In separate blocks, participants either judged the observed movement (task-relevant condition) or the colour of a dot on the screen (task-irrelevant condition). **(B)** Expectation manipulation in the main task. The avatar movement on the screen could either be expected (matching) or unexpected (mismatching) based on the participant’s own action, each occurring on 50% of trials. **(C)** Decoding of observed movements in the no-movement trials. In separate no-movement blocks, participants observed index and little finger abductions on the screen without moving themselves and while performing a detection task at fixation (top). These blocks provided independent data to train our classifier (middle). When training and testing the classifier on the no-movement data from all electrodes, the temporal generalization matrix (bottom right) revealed a significant cluster of above-chance decoding (outlined in black), primarily along the diagonal, with stimulus onset at 0 ms and peak decoding at 160 ms. The time course above shows decoding accuracy across testing time averaged over training times from 140-180 ms (window centred on peak decoding) and served as the baseline for deriving the predicted time courses in (D). The topographical map (bottom left) shows the corresponding Haufe-transformed classifier weights. **(D)** Predicted decoding time courses under the different theoretical accounts. Bayesian accounts (top) predict higher decoding for expected than unexpected outcomes. If implemented via pre-activation, this advantage should be evident already before stimulus onset (0 ms). Cancellation accounts (middle) predict higher decoding for unexpected outcomes, reflecting attenuation of expected inputs (or, equivalently, enhancement of unexpected inputs). The opposing process theory (bottom) predicts a reversal across time: higher decoding for expected outcomes early in time due to pre-activation, followed by later enhancement of unexpected outcomes. Columns with higher contrast reflect stronger sensory representations (high signal-to-noise ratio). See Supplementary Materials for details about how these predicted time courses were derived.

## Results

In the main task blocks, participants (*N* = 36) abducted either their index or little finger in response to a shape cue and observed a synchronous outcome on the screen: an avatar hand performing either the expected (matching) or unexpected (mismatching) movement relative to their own action. Expected and unexpected trials were intermixed within each block, and each occurred on 50% of trials (Figure 1B). The manipulation thus relied on expectations built up over participants’ lifetime of experience regarding the outcomes of action. These expectations are thought to be highly precise and to generate large error signals when violated (38,39), making this paradigm a good test of the opposing process theory. The task relevance of the observed movement was manipulated between blocks: In task-relevant blocks, participants judged the observed movement (e.g., ‘Did the index finger move?’), whereas in task-irrelevant blocks, they judged the colour of a dot presented on the screen (e.g., ‘Was the dot blue?’; Figure 1A). These main task blocks were followed by no-movement blocks, where participants passively observed index and little finger abductions on the screen while performing a detection task at fixation, providing independent data to train our pattern classifier (Figure 1C). At the end of the study, participants also provided surprise and probability ratings for expected and unexpected outcomes. The study was pre-registered at *https://doi.org/10.17605/OSF.IO/ZXR97*.

### Behavioural results

When the observed outcomes were task-relevant, participants were more accurate in responding to expected compared to unexpected outcomes (expected: *M* = 95.73%, *SD* = 3.61%; unexpected: *M* = 94.87%, *SD* = 4.00%; *t*(35) = 2.40, *p* =.022; Figure 2A). Reaction times were faster for unexpected than expected outcomes irrespective of task relevance, as reflected in a main effect of expectation (expected: *M* = 693.49 ms, *SD* = 105.02 ms; unexpected: *M* = 684.24 ms, *SD* = 102.11 ms; *F*(1, 35) = 5.64, *p* =.023; Figure 2B). Participants rated unexpected action outcomes as more surprising on a 0-100 scale (expected: *M* = 20.58, *SD* = 21.07; unexpected: *M* = 52.43, *SD* = 30.01; *t*(35) = -5.93, *p* <.001) and less probable (expected: *M* = 58.63, *SD* = 12.29; unexpected: *M* = 44.75, *SD* = 13.01; *t*(35) = 3.57, *p* =.001; Figure 2C). Taken together, these differences between expected and unexpected trials indicate that our expectation manipulation was effective. In the no-movement blocks, detection performance also indicated that participants were successfully engaged by the task (hit rate: *M* = 91.78%, *SD* = 10.13%; false alarm rate: *M* = 1.57%, *SD* = 5.00%). See Supplementary Materials for full analysis of the main task behavioural data.

**Figure 2:**
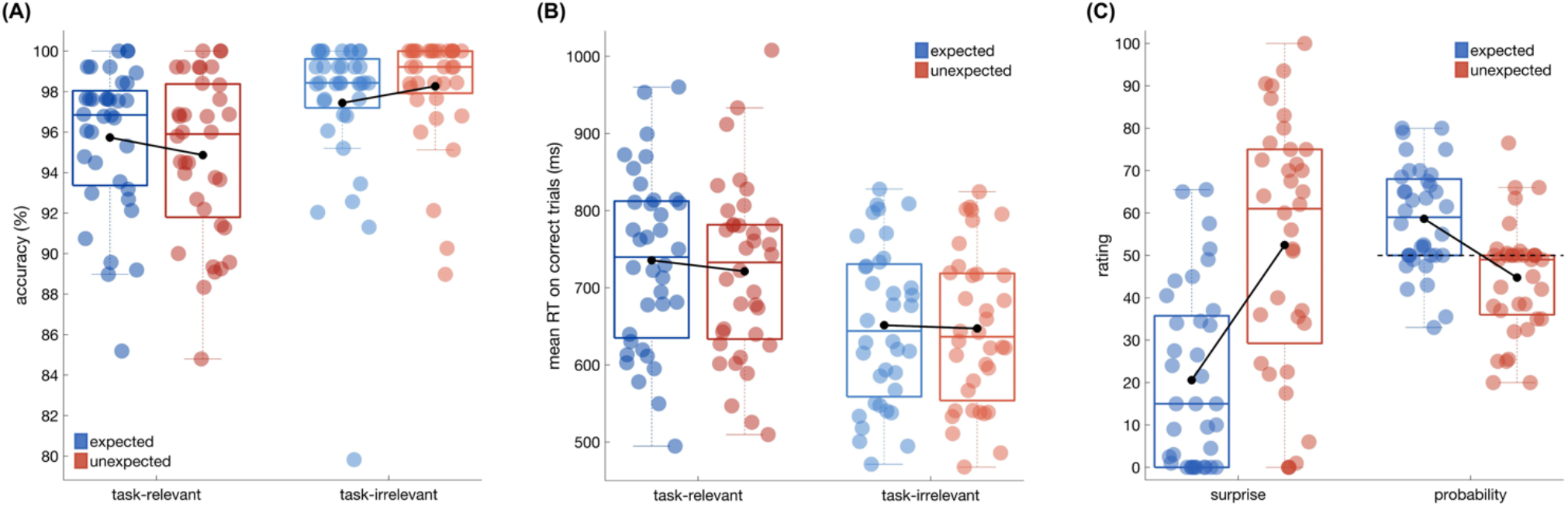
Behavioural and rating results. **(A)** Accuracy for expected and unexpected outcomes, separately for task-relevant and -irrelevant conditions. Accuracy was higher for expected than unexpected outcomes, when the expectations were task-relevant, whereas no expectation effect was observed when they were task-irrelevant. **(B)** Reaction times for expected and unexpected outcomes, showing faster responses to unexpected outcomes irrespective of task relevance. **(C)** Surprise and probability ratings for expected and unexpected outcomes, with expected outcomes rated as less surprising and more probable than unexpected outcomes. Boxes indicate the lower, middle, and upper quartiles, and whiskers extend to 1.5 times the interquartile range. Coloured dots represent individual participants, and black dots denote the group mean.

### EEG results – Decoding of observed movements in the no-movement blocks

We first examined whether observed finger movements could be decoded from EEG signals in the no-movement blocks. A linear classifier could indeed reliably distinguish observed index and little finger movements when trained and tested on the EEG data from all electrodes using cross-validation. The resulting temporal generalization matrix revealed a significant cluster of above-chance decoding (*p* <.001), primarily along the diagonal, spanning from around 100 to 300 ms after stimulus onset with peak decoding accuracy at 160 ms. The corresponding Haufe-transformed classifier weights were strongest over posterior electrodes and showed a lateralised pattern across the two hemispheres, consistent with sensitivity to the different spatial locations of the observed finger movements (Figure 1C).

### EEG results – Decoding of observed movements in the main task blocks

Our primary question was whether the influence of expectation on perceptual processing reversed across time, and whether any effects were modulated by task relevance. To address this question, we applied the classifier trained on the no-movement blocks to the main task data to decode the observed movements. For condition comparisons, decoding accuracy was averaged within the training time window in which decoding in the no-movement blocks peaked (140-180 ms), as pre-registered, yielding a one-dimensional decoding time course over testing time for each condition. When collapsing across task relevance, decoding accuracy differed between expected and unexpected outcomes (from approximately 135 to 240 ms; *p* =.003), whereas no overall difference was observed between task-relevant and -irrelevant outcomes (Figure S1A-D). Crucially, however, the effect of expectation differed as a function of task relevance (reflected in two significant clusters from approximately -80 to -35 ms [*p* =.042] and 20 to 70 ms [*p* =.019]; Figure S1E-F), motivating separate comparisons of expected and unexpected outcomes within each task. When outcomes were task-relevant, decoding accuracy was higher for expected than unexpected outcomes prior to outcome onset, corresponding to a significant cluster from approximately -95 to -10 ms (*p* =.015). This pattern reversed later in time, with decoding significantly higher for unexpected than expected outcomes from approximately 155 to 235 ms after outcome onset (*p* =.004; Figure 3A-B). When outcomes were task-irrelevant, only this later effect was observed, with higher decoding for unexpected than expected outcomes from approximately 145 to 240 ms post-stimulus (*p* =.004; Figure 3C-D). Together, these results demonstrate a rapid reversal in neural prioritisation, with enhanced decoding of task-relevant expected outcomes before stimulus onset, followed by an advantage for unexpected outcomes within 160 ms of stimulus presentation. See Supplementary Materials for more information concerning the EEG analyses.

**Figure 3:**
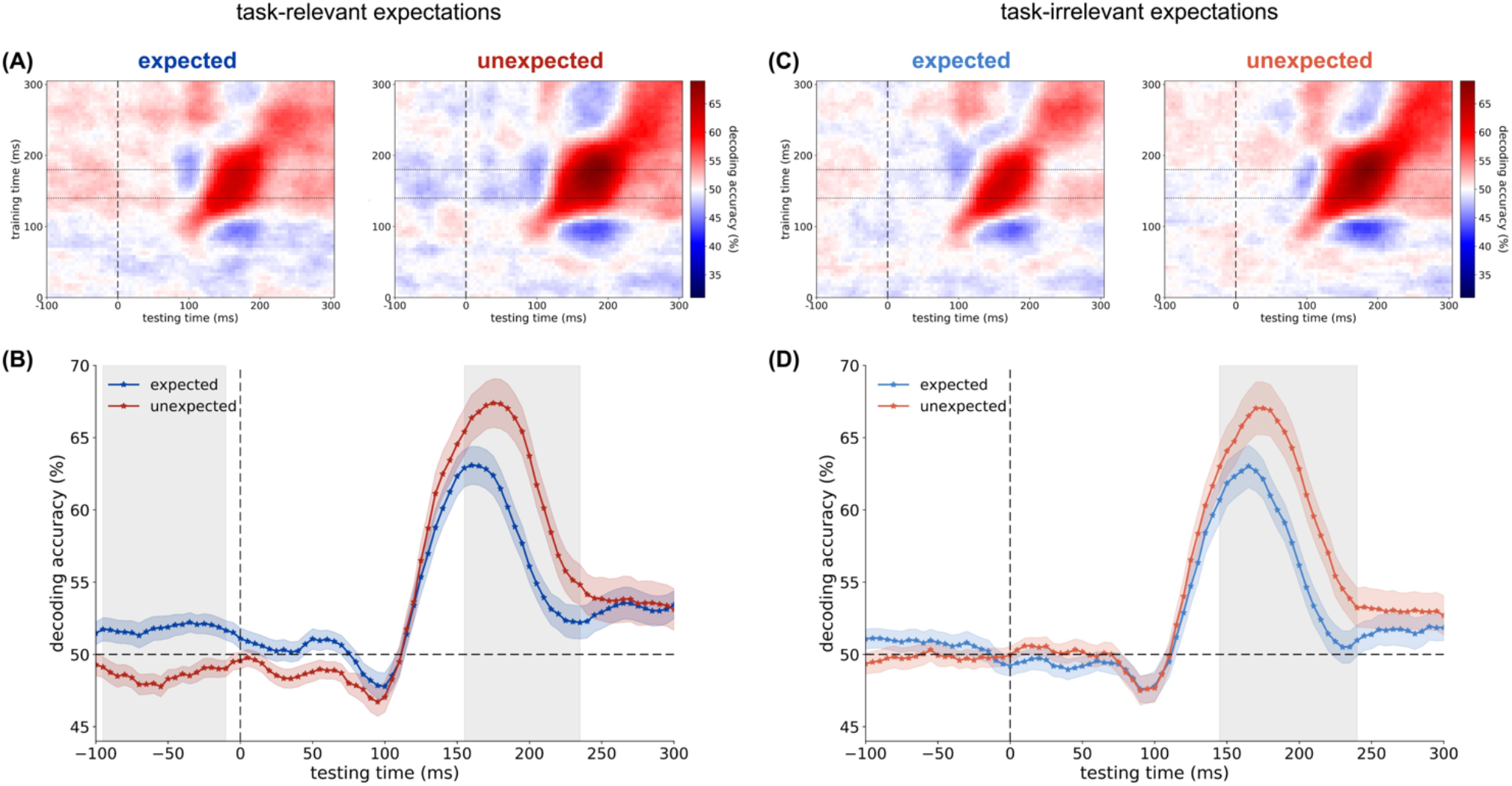
Main task decoding results. **(A)** Temporal generalization matrices for expected and unexpected outcomes in the task-relevant condition, with 0 ms marking the onset of the abducted hand image in both the no-movement (training) and main task (testing) trials. **(B)** Decoding time courses in the task-relevant condition, averaged across training times from 140 to 180 ms (centred on the decoding peak in the no-movement blocks). Decoding accuracy was higher for expected than unexpected outcomes prior to outcome onset (∼-95 to -10 ms). This pattern reversed after stimulus onset, with higher decoding for unexpected trials (∼155 to 235 ms). **(C)** Temporal generalization matrices for expected and unexpected outcomes in the task-irrelevant condition. **(D)** Decoding time courses in the task-irrelevant condition, showing significantly higher decoding for unexpected outcomes from approximately 145 to 240 ms. In the temporal generalization matrices, dotted horizontal lines mark the training time window used for averaging. In the time course plots, shaded regions represent the standard error of the mean (SEM), grey backgrounds indicate significant differences between expected and unexpected outcomes, and dashed horizontal lines mark chance level.

## Discussion

In this study, we observed a rapid temporal reversal in the neural prioritisation of expected relative to unexpected sensory events – specifically, observed action outcomes. We found higher decoding for task-relevant expected events starting before stimulus onset, followed by a later post-stimulus advantage for unexpected events within 160 ms. These findings are broadly in line with the opposing process theory and its predicted temporal dynamics, which aims to explain how expectations can render perception both veridical and informative. This theory builds upon the logic of current Bayesian and cancellation accounts but overcomes their limitation that prioritising a single adaptive advantage comes at the direct cost to another. Instead, it is proposed that through these temporal dynamics, we can rapidly generate largely veridical percepts, while remaining sensitive to highly surprising events that are informative for learning and model updating.

This rapid reversal in prioritisation may help to reconcile previously conflicting findings concerning the typical effects of expectation on perception and neural responses (13,27,29–31,33), as well as mixed findings reported in clinical populations (40,41). In behavioural paradigms, perception is of course typically probed with a single response per trial (e.g., 18,27), such that either the early or later process may dominate the resulting behavioural measure. According to the opposing process theory, the ‘winning’ process depends on the magnitude of the error signal and the timing of perceptual judgements (36,42). Likewise, functional magnetic resonance imaging (fMRI) lacks the temporal resolution necessary to dissociate these rapidly unfolding effects, such that the measured signal likely reflects a mixture of both processes. However, its superior spatial resolution has also revealed interesting dissociations across cortical layers (43), which could be related to these temporal dynamics in future work.

Interestingly, even in electrophysiological studies with sufficient temporal resolution, these dynamics are not typically observed. We believe that this is largely due to differences in paradigms and analysis decisions. In the present study, the advantage for expected events emerged before stimulus onset, in line with proposals of predictive pre-activation, and common analysis choices can obscure such effects. Specifically, studies examining predictive processing of sensory events often baseline-correct to immediately before stimulus onset (44,45) which effectively removes any biases that begin pre-stimulus. Notably, this early effect was restricted to our task-relevant condition, although it remains unclear whether it was entirely absent or merely weaker when outcomes were task-irrelevant (see also 31,34,35,46,47). Importantly, such analysis norms for baseline correction also complicate the interpretation of post-stimulus benefits observed for unexpected inputs. Given the pre-activation in the present study, apparent enhancements of unexpected outcomes may instead often reflect subtracted pre-activation of the expected. In contrast, the present findings suggest a genuine later enhancement of unexpected inputs, as it emerges alongside the pre-activation of expected outcomes. Other studies that avoid such baseline-correction choices also do not necessarily find the later enhancement for unexpected inputs. This may be because, under the opposing process theory, the later effect is proposed to only occur for highly surprising events (defined according to Kullback-Leibler Divergence; 36) that are informative for learning. Most previous studies use ‘unexpected’ conditions that would not generate such high surprise – for example, when expectations are learned within the study and violations fall within the learned variability of the inputs (e.g., 31), constituting ‘expected uncertainty’ (48). In contrast, the unexpected events in the present study violated expectations built across a lifetime of learned highly precise priors about the outcomes of action (38,39), and hence provided a good test for the predictions of the opposing process theory.

The precise mechanism underlying the later enhancement observed for our unexpected outcomes remains unclear. One possibility is that it is driven by phasic catecholamine release in response to surprising events, as suggested by (36). Surprising events are known to trigger such phasic neuromodulatory responses (49,50), and tonic increases in noradrenaline have been shown to amplify the signal-to-noise ratio in sensory systems (51). It is therefore plausible that such a reactive gain control mechanism generates the unexpected enhancement observed here (see also 52). For this mechanism to explain the findings, it would likely require that the action predictions are relatively ‘stubborn’ – that is, not updated in response to the unexpected outcomes and continuing to elicit processes typically associated with ‘unexpected uncertainty’ (50). We predicted this would be the case given the typical precision of action predictions (38), hence the present study design. Future work could test this possibility more directly, for example via pharmacological intervention or by analysing pupillary responses in these paradigms. The unexpected enhancement observed here could also arise from other mechanisms, including suppression of expected inputs, as proposed by cancellation accounts. Importantly, however, the observed temporal reversal in neural prioritisation is uniquely predicted by the opposing process theory, and neither Bayesian nor cancellation accounts predict such a dynamic.

We here studied the influence of expectations on perception in an action context. We thus exploit the high precision of action expectations (38), which is one way to generate large error signals upon presentation of unexpected sensory events. It is these larger error signals which we believe may be a reason that the cancellation logic has prevailed in action disciplines across the decades – for example, thought to explain why we cannot tickle ourselves and why expected action outcomes feel less intense (22). However, much work from the last decade has suggested that the underlying mechanisms via which expectation shapes perception may be broadly similar across active and passive domains of sensory processing (11,18,34,36,53), and thus we would expect our findings to generalise beyond action contexts. Some early support for this idea comes from a recent study reporting enhanced decoding of unexpected scene categories alongside earlier expected enhancements (54), although notably, all effects in that study were later and post-stimulus, perhaps due to the higher order categories decoded.

To conclude, we observed a temporal reversal in the neural prioritisation of expected relative to unexpected sensory events, with higher decoding for task-relevant expected events before stimulus onset, consistent with pre-activation of the expected sensory representation, followed by a later post-stimulus advantage for unexpected events within 160 ms. These findings are broadly in line with the opposing process theory and its predicted temporal dynamics, which may explain how expectations can render perception veridical, while still allowing for the accurate perception of particularly unexpected events that are important for learning. These findings may thus help reconcile previously conflicting results and accounts, with relevance for theories of both learning and perception across sensory domains.

## Methods

### Participants

Thirty-six participants (mean age = 27.69 years, *SD* = 7.34; age information missing for one participant) completed the study and were included in the analysis. An additional two participants were excluded from the analysis because their behavioural accuracy on unexpected trials in the task-relevant condition was below 3%, suggesting that they misunderstood the task – appearing to respond based on the finger they themselves moved rather than the finger moving on the screen. All participants had normal or corrected-to-normal vision. The study complied with all relevant ethical regulations and was approved by the Research Ethics Committee at Birkbeck, University of London. Participants gave informed consent prior to their participation and either received course credits or £25, equivalent to an hourly wage of approximately £8.

### Stimuli

We used the same stimuli as in (34), displaying a gender-neutral right hand viewed from a canonical first-person perspective. This hand subtended approximately 15° in height and 8° in width of visual angle. A white fixation cross was centrally placed on the middle finger and could be surrounded by either a circle or a square. Coloured dots (red or blue) were positioned at the location where the fingertip of the abducted finger had been prior to movement. Stimuli were presented against a black background on a 24-inch monitor (1920 x 1080 pixel resolution, 60 Hz refresh rate), with the base of the hand aligned to emerge from the lower edge of the screen. Participants rested their own right hand on a support aligned with the monitor’s lower edge. Their hand was rotated approximately 45° counterclockwise from the vertical midline (i.e., 12 o’clock) and was concealed from view by a black box to prevent visual feedback from their movements.

### Procedure

The experiment was programmed in MATLAB using Psychtoolbox (55) and closely followed the design of a previous fMRI study (34). On each trial, participants performed a movement (i.e., abducting their index or little finger) and were either presented with the expected or unexpected action outcome on the screen, i.e., an avatar hand performing an abduction movement either matching or mismatching the participants’ own action, and presented in synchrony with their movement.

Each trial began with a fixation cross displayed for 500 ms and until participants pressed down two keys on a number pad with the index and little finger of their right hand. Then, a neutral hand image appeared on the screen. After 500 ms, and provided both keys were still pressed, a shape (circle or square) appeared around the fixation cross, instructing participants whether to abduct their index or little finger. The shape-action mapping reversed halfway through the main task, and the first mapping was counterbalanced across participants. As soon as participants performed the correct movement, the neutral hand image was replaced by an image of the avatar hand abducting either its index or little finger, creating apparent motion. On 50% of trials, the avatar’s movement matched the participant’s own action, and on the other 50% it did not. The movement also revealed a coloured dot (red or blue) at the previous fingertip location. The hand image was removed after 500 ms, followed by a blank screen for 500 ms (250 ms for the first two participants).

This blank screen was followed by a 1500 ms response window with participants performing one of two tasks: They either judged the observed finger abduction (e.g., ‘Did the index finger move?’; task-relevant condition) or the colour of the revealed dot (e.g., ‘Was the dot blue?’; task-irrelevant condition). The task was blocked and alternated across blocks, with the first task counterbalanced across participants. Responses (‘yes’/’no’) were made with the left thumb on a separate number pad. Each trial ended with a random inter-trial interval (ITI) between 1000 and 1500 ms. Participants completed four blocks of each task, with 64 trials per block. Trial order within each block was randomized, and participants received feedback about their accuracy and mean reaction time at the end of each block. Prior to this, participants completed one block with 40 trials for each task as practice, during which the outcome always matched their own action.

After the eight main task blocks, participants completed three no-movement blocks, each with 80 trials. During these blocks, participants were shown index and little finger abductions on the screen in a random order, without performing any movements themselves and while doing a task at fixation. The data from the no-movement blocks were used in the analysis to train a classifier to discriminate between the two observed action outcomes. Each no-movement trial began with a fixation cross shown for 500 ms, followed by the neutral hand image for 1000 ms (without a shape cue around fixation). This was then replaced by an avatar hand abducting either its index or little finger, shown for 500 ms. Unlike in the main task, no coloured dot appeared under the abducted fingertip. The trial ended with a random inter-trial interval between 1000 and 1500 ms. During these no-movement blocks, participants performed a detection task at fixation: On 10% of trials, the fixation cross was absent for the first 25 ms of the abducted avatar hand presentation, and participants had to press a key when they detected this. They received feedback concerning their hit and false alarm rates after each block.

At the end of the study, participants were asked to rate how surprised they felt by the different outcomes, as well as their probability (e.g., ‘How surprised were you when you lifted your index finger and the little finger on the screen moved?’ and ‘When you lifted your index finger, how often did the little finger on the screen move?’) on a scale from 0 (‘not surprised at all’/’never (0%)’) to 100 (‘highly surprised’/’always (100%)’).

### EEG recording and preprocessing

Continuous EEG data were DC-recorded using the BrainVision recorder software and a BrainAmp DC amplifier (Brain Products, Munich, Germany). Sampling rate was 500 Hz with a resolution of 0.1 μV. The upper cut-off frequency during DC recording was 250 Hz, and EEG was digitally low pass filtered at 40 Hz. EEG was recorded from 27 scalp sites at Fpz, F7, F3, Fz, F4, F8, FC5, FC6, T7, C3, Cz, C4, T8, CP5, CP6, P9, P7, P3, Pz, P4, P8, P10, PO9, PO7, PO8, PO10, and Oz. All electrodes were online referenced to the left earlobe and impedances were kept below 5 kΩ.

Preprocessing was conducted in MATLAB using FieldTrip (56). The continuous EEG data were first re-referenced to the average of both earlobes. Only to identify ICA components, the data were band-pass filtered between 1 and 40 Hz using a FIRWS filter. Artifact components, such as those reflecting heartbeats or eye blinks, were then identified and removed from the unfiltered data. Next, the data were segmented into trials and these were visually inspected for artifacts. To aid artifact detection, the variance (collapsed over channels and time) was calculated for each trial. Trials with large variance were selected for manual inspection and removed if they contained excessive or irregular artifacts. Finally, the data were baseline-corrected using the 100 ms interval before cue onset for the main task (i.e., the shape cue around fixation) and before the onset of the abducted hand image for the no-movement trials.

### Behavioural data analysis

For the main task, all trials with RTs below 100 ms were excluded from the analysis. Additionally, one block of the task-relevant condition was excluded for three participants: One had responded in the first block based on which finger they moved rather than which action outcome was shown on the screen, and two had used the wrong response keys for one block, resulting in systematically incorrect responses. For accuracy and RTs for correct trials, we conducted 2×2 repeated-measures ANOVAs with the factors expectation (expected, unexpected) and task relevance (task-relevant, task-irrelevant). In cases of a significant interaction, we followed up with comparisons of expected versus unexpected trials within each task.

For participants’ surprise and probability ratings, we compared the mean ratings for expected and unexpected trials using paired-samples t-tests. For the no-movement blocks, key presses occurring between 100 and 1500 ms after the onset of the abducted hand image (and the onset of the absent fixation cross) were considered detection responses. We then calculated participants’ hit and false alarm rates.

### Decoding within the no-movement blocks

No-movement trials in which the fixation cross disappeared or participants made a detection response were excluded from the analysis. Decoding was performed in Python using scikit-learn (57), with linear discriminant analysis (LDA) as the classifier. The shrinkage parameter was set to ‘auto’ and the solver to ‘eigen’, which ensures that the covariance of the data is taken into account. A two-way classifier was trained on the no-movement data from all 27 electrodes to discriminate between the two action outcomes (i.e., index vs. little finger abduction) and tested on the same data using a cross-validation approach. To ensure balanced training data, the no-movement trials were randomly downsampled so that there was the same number of trials for each action outcome after artifact rejection. The classifier was then trained on all trials except two (one from each class) and tested on these left-out trials. This procedure was repeated until all trials had been left out once. Training and testing were performed in a time-resolved manner, yielding a training-by-testing time (temporal generalization) matrix. The classifier was trained at each time point from 0 to 300 ms after the onset of the abducted hand image and tested from -100 to 300 ms relative to onset, in steps of 5 ms. For each time point, the data from each electrode were averaged within a 30 ms window centred on the time point of interest. The training data from each electrode were z-scored across trials, and the testing data were z-scored using the mean and standard deviation from the training data.

To assess whether decoding accuracy in the temporal generalization matrix differed significantly from chance (i.e., 50%), we used cluster-based permutation tests to identify temporal clusters (58). For the two-dimensional clusters, elements were considered neighbours if they were directly adjacent, either cardinally or diagonally. The cluster-forming threshold corresponded to a p-value below.05 (two-tailed). Cluster-level statistics were computed as the sum of t-values within each cluster and compared to a null distribution generated from 10,000 random permutations. Clusters were considered significant if their p-value was below.05 (two-tailed).

To examine which electrodes contributed to the classifier’s discrimination of observed index versus little finger abductions, we computed Haufe-transformed activation patterns (59) from the LDA weights. For each training time point, the classifier weights were multiplied by the covariance of the unstandardized training data, yielding forward-model patterns in electrode space. The resulting patterns were averaged across cross-validation folds and participants, and visualised as topographical maps.

### Main task decoding

We applied the classifier trained on the no-movement data to the main task to decode the observed action outcomes, separately for expected and unexpected trials, task-relevant and -irrelevant blocks, as well as their combinations. Training and testing were again performed in a time-resolved manner in steps of 5 ms, using the same training and testing time ranges as for the decoding within the no-movement blocks, as well as the same data averaging and z-scoring procedures at each time point. This resulted in a temporal generalization matrix for each condition.

To compare conditions, we averaged decoding accuracy across training times from 140 to 180 ms (a window centred on the decoding peak along the diagonal for the no-movement blocks, following 31). This generated a one-dimensional time course over testing time (from -100 to 300 ms relative to action outcome onset) for each condition. To test for condition differences in decoding, we again performed cluster-based permutation tests comparing the relevant paired conditions. All cluster test parameters were identical to those described earlier.

For all decoding analyses, we excluded the first task-relevant block of the participant who had responded based on their own movement rather than the observed action outcome in that block.

## Acknowledgements

This work was funded by a European Research Council (ERC) consolidator grant (101001592) under the European Union’s Horizon 2020 research and innovation programme, awarded to CP. We are grateful to Rebecca Nako for training and assistance with the EEG.

