## Supplementary Materials for "Paradoxical influences of prediction are resolved across time"

### Paradoxical influences of prediction are resolved across time – Supplementary Materials

#### Behavioural results

We here report the results of the full 2x2 ANOVAs, concerning how expectation (expected vs. unexpected) and task relevance (relevant vs. irrelevant) impact upon accuracy and reaction time.

For accuracy, there was no main effect of expectation (expected: $M$ = 96.59%, $SD$ = 3.49%; unexpected: $M$ = 96.57%, $SD$ = 3.13%; $F(1, 35) = 0.004$, $p = .951$). There was, however, a significant main effect of task relevance ($F(1, 35) = 43.92$, $p < .001$), with higher accuracy in the task-irrelevant (colour task; $M$ = 97.85%, $SD$ = 3.05%) than in the task-relevant (finger task; $M$ = 95.30%, $SD$ = 3.65%) blocks. Importantly, these two factors interacted ($F(1, 35) = 14.07$, $p < .001$). To follow up this interaction, we examined the effect of expectation separately for each task. When expectations were task-relevant, participants responded more accurately to expected than unexpected outcomes (expected: $M$ = 95.73%, $SD$ = 3.61%; unexpected: $M$ = 94.87%, $SD$ = 4.00%; $t(35) = 2.40$, $p = .022$). In contrast, when expectations were task-irrelevant, accuracy did not significantly differ between expected and unexpected trials (expected: $M$ = 97.44%, $SD$ = 3.82%; unexpected: $M$ = 98.27%, $SD$ = 2.73%; $t(35) = -1.87$, $p = .070$).

For reaction times, there was a significant main effect of expectation ($F(1, 35) = 5.64$, $p = .023$), with faster responses to unexpected ($M$ = 684.24 ms, $SD$ = 102.11 ms) than expected ($M$ = 693.49 ms, $SD$ = 105.02 ms) outcomes. There was also a significant main effect of task relevance ($F(1, 35) = 66.22$, $p < .001$), with faster responses in the task-irrelevant (colour task; $M$ = 649.30 ms, $SD$ = 100.90 ms) than in the task-relevant (finger task; $M$ = 728.43 ms, $SD$ = 112.72 ms) blocks. The interaction between expectation and task relevance was not significant ($F(1, 35) = 3.91$, $p = .056$).

#### EEG results – Decoding of observed movements in the main task blocks

When collapsing across task relevance, decoding accuracy differed between expected and unexpected outcomes, with higher decoding for unexpected trials from approximately 135 to 240 ms (*p* = .003; Figure S1A-B). In contrast, collapsing across expectation revealed no difference in decoding accuracy between the task-relevant and -irrelevant condition (Figure S1C-D). Importantly, the effect of expectation differed as a function of task relevance, reflected in two significant clusters from approximately -80 to -35 ms (*p* = .042) and 20 to 70 ms (*p* = .019; Figure S1E-F). Across both clusters, the expectation effect (expected – unexpected) was more positive in the task-relevant than the -irrelevant condition.

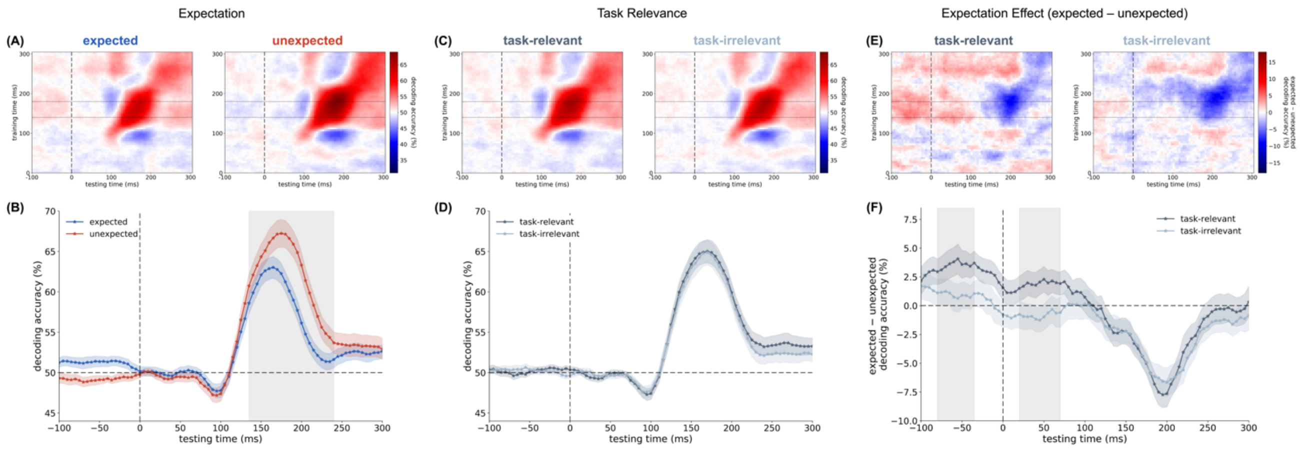


**Figure S1: Main task decoding results. (A)** Temporal generalization matrices for expected and unexpected outcomes, collapsed across task relevance. **(B)** Decoding time courses collapsed across task relevance, showing higher decoding for unexpected than expected outcomes (around 135 to 240 ms). **(C)** Temporal generalization matrices for task-relevant and -irrelevant outcomes, collapsed across expectation. **(D)** Decoding time courses collapsed across expectation, showing no difference between task-relevant and -irrelevant outcomes. **(E)** Temporal generalization matrices showing the expectation effect (expected – unexpected) separately for task-relevant and -irrelevant outcomes. **(F)** Expected minus unexpected decoding time courses by task relevance, showing a more positive expectation effect in the task-relevant than -irrelevant condition (around -80 to -35 ms and 20 to 70 ms). In the temporal generalization matrices, 0 ms denotes the onset of the abducted hand image in both the no-movement (training) and main task (testing) trials. Dotted horizontal lines mark the training time window used for averaging (140 to 180 ms). In the time course plots, shaded regions indicate SEM, grey backgrounds mark significant differences between conditions, and dashed horizontal lines indicate chance level (B, D) or zero (F).

#### EEG results – ERPs

We additionally examined whether univariate event-related potentials (ERPs) in the main task differed as a function of expectation and task relevance. The same data were included as in the main task decoding analysis. ERPs were computed from -100 ms relative to the onset of the abducted hand image until its offset at 500 ms. To assess effects of expectation and task relevance, we used cluster-based permutation tests to identify spatiotemporal clusters across electrodes and time points. All cluster test parameters were identical to those used in the decoding analyses.

When collapsing across task relevance, ERPs differed between expected and unexpected outcomes. One significant cluster showed a more negative response for unexpected compared to expected trials (*p* = .007; approximately 190 to 275 ms), followed by a second cluster showing a more positive response for unexpected trials (*p* < .001; approximately 270 to 500 ms). Both clusters extended over nearly all electrodes (Figure S2A). When collapsing across expectation, ERPs also differed as a function of task relevance. A significant cluster indicated a more positive response in the task-irrelevant than the -relevant condition ($p$ = .011; approximately 270 to 500 ms), predominantly over posterior electrodes (Figure S2B). The effect of expectation also varied with task relevance, as reflected in a significant cluster from approximately 170 to 250 ms ($p$ = .026), extending over most electrodes (Figure S2C). This cluster indicated a larger difference between expected and unexpected outcomes in the task-relevant condition.

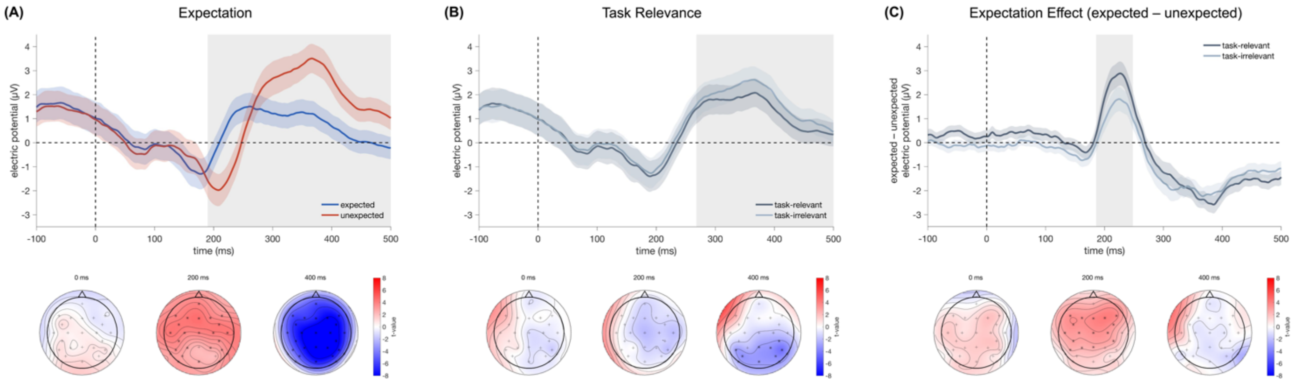


**Figure S2: ERP results. (A)** ERPs for expected and unexpected outcomes, collapsed across task relevance. Unexpected outcomes elicited a more negative response from approximately 190 to 275 ms, followed by a more positive response from around 270 to 500 ms, extending over most electrodes. **(B)** ERPs for task-relevant and -irrelevant conditions, collapsed across expectation. Responses were more positive in the task-irrelevant than the -relevant condition from around 270 to 500 ms, predominantly over posterior electrodes. **(C)** Expectation effect (expected – unexpected) shown for the task-relevant and -irrelevant conditions. A significant cluster was observed from around 170 to 250 ms over most electrodes, reflecting a larger difference between expected and unexpected outcomes in the task-relevant condition. Time-course plots show ERPs averaged across all electrodes from 100 ms before stimulus onset until its offset at 500 ms. Shaded regions indicate the SEM, and grey backgrounds denote time windows during which any electrode was part of a significant cluster. Topographical maps show t-values used for cluster formation at 0, 200, and 400 ms post-stimulus onset, with electrodes belonging to a significant cluster marked with stars and all others with crosses.

Given that the effect of expectation varied with task relevance, we next compared ERPs for expected and unexpected outcomes separately for the task-relevant and -irrelevant conditions. In the task-relevant condition, unexpected outcomes elicited a more negative response from approximately 190 to 275 ms ($p$ = .004), followed by a more positive response from approximately 270 to 500 ms ($p$ < .001; Figure S3A), both extending across nearly all electrodes. In the task-irrelevant condition, ERPs also differed between expected and unexpected outcomes: Unexpected outcomes again elicited a more negative response from approximately 195 to 265 ms ($p$ = .016), followed by a more positive response from approximately 270 to 500 ms ($p$ < .001; Figure S3B), spanning most electrodes.

Univariate effects like these are sometimes taken to support cancellation accounts, as they indicate larger overall responses to unexpected than expected outcomes. However, such effects only tell us about the magnitude of overall processing and not the content of activation. In contrast, the main decoding analyses of interest pertain to multivariate differences between index and little finger representations within each condition. Whereas the ERPs collapse across finger identity and capture overall condition-averaged responses, the decoding analyses assess stimulus-specific information.

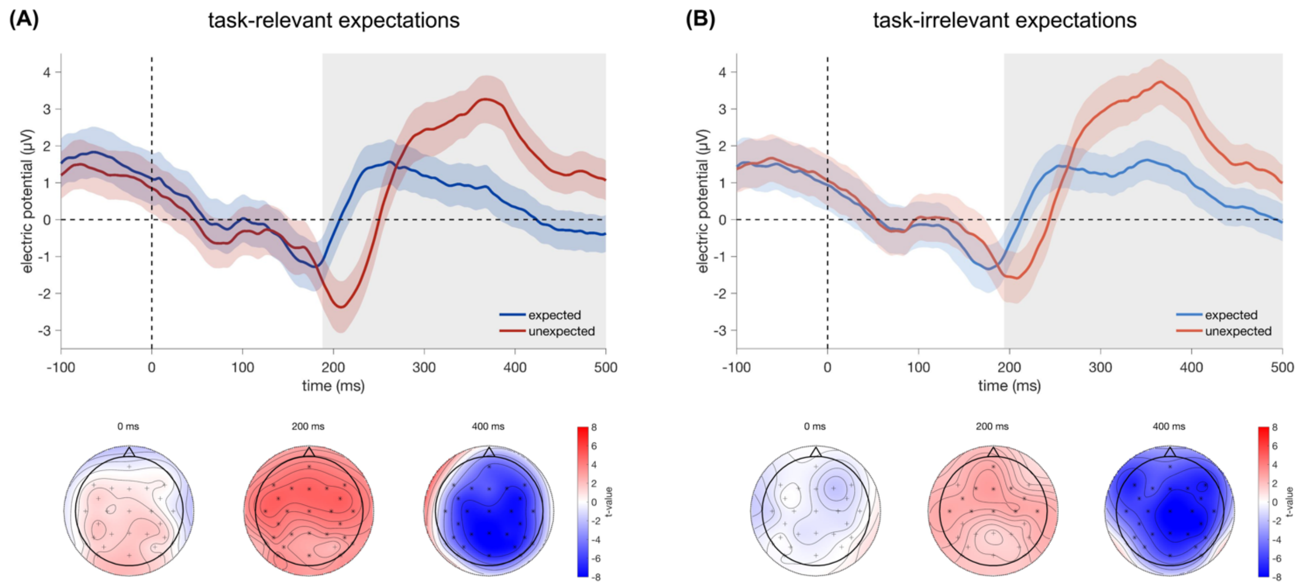


**Figure S3: ERP results within task-relevant and -irrelevant conditions. (A)** ERPs for expected and unexpected outcomes in the task-relevant condition. Unexpected outcomes elicited a more negative response from around 190 to 275 ms, followed by a more positive response from around 270 to 500 ms, extending across nearly all electrodes. **(B)** ERPs for expected and unexpected outcomes in the task-irrelevant condition. Unexpected outcomes similarly elicited a more negative response from around 195 to 265 ms, followed by a more positive response from around 270 to 500 ms, also across most electrodes. Time-course plots show ERPs averaged across all electrodes from 100 ms before stimulus onset to its offset at 500 ms. Shaded regions indicate the SEM, and grey backgrounds denote time windows during which any electrode was part of a significant cluster. Topographical maps show t-values used for cluster formation at 0, 200, and 400 ms post-stimulus onset, with electrodes belonging to a significant cluster marked with stars and all others with crosses.

**Predicted decoding time courses in Figure 1**

To derive predicted decoding time courses under the different theoretical accounts in Figure 1, we used the decoding time course from the no-movement blocks as a baseline. This time course reflects decoding of the observed actions in the absence of an expectation manipulation and served as a neutral baseline onto which the predicted expectation effects were applied. Specifically, decoding accuracy was averaged across training times from 140 to 180 ms (a window centred on the peak decoding along the diagonal), yielding a one-dimensional time course over testing time, mirroring the analysis approach used for condition comparisons in the main task.

Under Bayesian accounts, expectations are proposed to enhance the neural representations of expected outcomes. We operationalised this enhancement as pre-activation of the expected sensory representations. Specifically, we implemented this as an additive modulation of the decoding signal. For expected trials, this modulation corresponds to sensory evidence consistent with the subsequently presented outcome, whereas for unexpected trials it corresponds to sensory evidence for the alternative outcome. Accordingly, a constant offset was added to the baseline decoding time course for expected trials and subtracted for unexpected trials. This modulation amounted to ±2% decoding accuracy applied uniformly across all time points, yielding differences between the predicted expected and unexpected time courses similar to those reported previously (1). We did not have particular predictions about effect size and ±2% was chosen for illustrative purposes in Figure 1.

Under cancellation accounts, expected inputs are proposed to be suppressed, such that unexpected outcomes are relatively prioritised. For our predicted time courses in Figure 1, we operationalised this mechanism as a selective enhancement of unexpected outcomes rather than explicit suppression of expected ones, but these two implementations are equivalent in terms of the resulting differences between expected and unexpected decoding time courses. Starting from the baseline decoding time course, we added a time-varying enhancement to the decoding signal for unexpected trials, while leaving the expected time course unchanged. This enhancement was implemented as a smooth increase in decoding accuracy anchored to the time at which baseline decoding first rose above chance (115 ms post-stimulus onset, as determined by a cluster-based permutation test against chance). The enhancement ramped up smoothly around this time point (with an exponential time constant of 20 ms), reaching a maximum amplitude of 7% decoding accuracy that was then sustained across the remainder of the post-stimulus period. This resulted in higher decoding for unexpected relative to expected outcomes once stimulus information was available, resembling patterns reported in previous work (2).

The opposing process theory proposes early enhancement of expected outcomes followed by later prioritisation of unexpected outcomes. We modelled this account by combining the implementations used for the Bayesian and cancellation accounts. Specifically, the predicted time course for expected trials included the same constant pre-activation offset (2%) as in the Bayesian account, while the predicted time course for unexpected trials combined a reduction due to incorrect pre-activation with later enhancement as implemented for the cancellation account. The amplitude of the post-stimulus enhancement was increased to 11% decoding accuracy (instead of 7%) to compensate for the reduction due to incorrect pre-activation, ensuring that the resulting late differences between expected and unexpected outcomes matched in magnitude those predicted under the cancellation account. Together, this implementation yielded an early advantage for expected outcomes followed by a later advantage for unexpected outcomes once outcome information was available.

**Deviations from the pre-registration**

There were a few minor deviations between the pre-registered design and the final implementation of the study. After collecting data from a few pilot participants, we noticed that the design was too fast with overlapping EEG responses to successive events. To address this, the timing of the trials was slowed. In the main task, the neutral hand image was presented for 500 ms before the cue appeared around fixation, rather than the hand and cue appearing simultaneously. In addition, the duration of the blank screen preceding the response was increased from 250 to 500 ms, and the ITI was extended to vary between 1000 and 1500 ms instead of being fixed at 500 or 1000 ms. Because these changes increased trial duration, the number of trials per block was reduced from 80 to 64 trials. The no-movement blocks were similarly slowed, with the neutral hand image presented for 1000 instead of 500 ms and the ITI increased to vary between 1000 and 1500 ms. As a consequence of the longer ITI, key presses occurring up to 1500 ms (rather than 1000 ms) after onset of the abducted hand image, so until the start of the next trial, were considered valid detection responses. Given the longer trial duration, the no-movement trials were split into three blocks of 80 trials instead of two blocks of 120 trials.

For preprocessing, we also decided to adopt a more minimal approach than pre-registered following reading of recommendations to reduce the possibility of artificially smearing our effects across time (3). Rather than bandpass filtering the data between 0.2 and 40 Hz, all analyses were conducted on the unfiltered data. Filtering was only applied to identify artifact-related ICA components, which were then removed from the unfiltered data. In addition, no detrending was applied. For baseline correction, we shortened the window to 100 ms.
